# The slit diaphragm in *Drosophila* features a bi-layered, fishnet-like architecture

**DOI:** 10.1101/2025.03.06.641875

**Authors:** Deborah Moser, Konrad Lang, Alexandra N. Birtasu, Margot P. Scheffer, Martin Helmstädter, Tobias Hermle, Achilleas S. Frangakis

## Abstract

The kidney filters large volumes of blood plasma, relying on the glomerulus for the filtration. The slit diaphragm, a critical component of the glomerulus, is formed between podocytes by the immunoglobulin domain proteins nephrin and Neph1. The molecular architecture of the slit diaphragm has remained elusive for decades. Using cryo-electron tomography on focused ion beam-milled *Drosophila* nephrocytes, an invertebrate podocyte model, we show that the slit diaphragm adopts a fishnet-like pattern. Comparison of hundreds of slit diaphragm segments reveals that it is bi-layered and highly periodic. Based on the cryo-electron tomography map, we propose four possible models for the arrangement of the *Drosophila* nephrin ortholog (Sns), and the *Drosophila* Neph1 ortholog (Kirre), the main components of the slit diaphragm in *Drosophila*. In each model, precise and consistent homo- and heterophilic interactions between crossing immunoglobulin domains of Sns and Kirre become apparent, with immediate implications for the stability and the assembly of the slit diaphragm. Cryo-electron tomography shows that *sns* silencing disrupts this fishnet pattern, linking this directly to *Drosophila* nephrin. After Rab5 silencing, causing Sns mistrafficking and ectopic SD formation, the fishnet pattern appears ectopically as well. Our findings align with observations applying cryo-electron tomography to podocytes in mice, indicating that the molecular architecture is evolutionarily conserved across animals. This highlights the value of the nephrocyte as a podocyte model and establishes a crucial link between the architecture of the slit diaphragm and its function.

## INTRODUCTION

To ensure essential clearance, the kidneys generate about 180 liters of primary urine daily. Retaining plasma proteins during filtration, the glomerular filter acts as a molecular sieve with three complementary layers: the fenestrated endothelium, the glomerular basement membrane and the slit diaphragm (SD), a specialized intercellular junction formed between the podocyte foot processes. The molecular framework of the SD is established by immunoglobulin(Ig)-domain-containing proteins nephrin^1–4^ and Neph1^5,6^, which engage extracellularly across the filtration slit^7,8^. These proteins assemble a multi-protein complex featuring a hybrid composition of junctional components^9,10^ to provide dynamic structural stability while integrating signaling cues^11,12^. DNA variants in *NPHS1*, the gene encoding nephrin, cause congenital nephrotic syndrome^1^, highlighting the slit diaphragm’s crucial role for proper kidney function.

Since its discovery in the mid-20th century, the molecular architecture of the SD has been a subject of ongoing debate^13^. Studying the SD is challenging, due to its absence from *in vitro* models, limited accessibility, and molecular complexity of the minute components. Using room temperature transmission electron microscopy (RT-TEM) on perfusion-fixed rodent kidneys, Rodewald and Karnovsky proposed a zipper-like structure^14^. Electron tomography showed molecular strands^15^ and a layered, bipartite arrangement^16^. Most recently, cryo-electron tomography (cryo-ET) of native murine glomeruli revealed that the SD has a fishnet-like pattern and proposed a molecular architecture of the SD^17^.

The podocyte-like nephrocyte in *Drosophila* is a genetically tractable model that enables mechanistic studies, variant analyses, and drug screening^18–21^. It emerged as a valuable model of the glomerular filter through its structural and molecular conservation^22–24^. Featuring an accessible SD, *Drosophila* is an attractive organism for investigating the molecular ultrastructure of the SD.

## SHORT METHODS

### Fly strains and husbandry

Flies were raised on standard *Drosophila* food at 25°C. *Prospero*-*GAL4*^22^ was used to control expression of *sns*-RNAi (Bloomington Drosophila Stock Center #64872) in garland cell nephrocytes, wild-type (yw^11^^18^) was crossed to *Prospero*-*GAL4* as control, further stocks are listed in **Supplemental Table 1**.

### Cryo-electron tomography

For cryo-electron tomography (cryo-ET), nephrocytes were dissected in Schneider’s medium containing 10% glycerol, centrifuged onto poly-L-lysin coated EM grids and plunge frozen in liquid ethane. Plunge frozen grids were imaged by cryogenic confocal laser scanning microscopy, then clipped into focused ion beam (FIB) autogrids. On-grid lamellae were FIB-milled in a Helios Nanolab 600i (Thermo Fisher Scientific Inc), and tilt series acquired in a Titan Krios transmission electron microscope (Thermo Fisher Scientific Inc.). Data was processed and subtomogram averaging performed using RELION-5^25^. Further details are described in the **Supplemental Methods**.

### Conventional transmission electron microscopy

For conventional transmission electron microscopy (TEM), nephrocytes were dissected in phosphate buffered saline (PBS) followed by fixation in a mix of 4% paraformaldehyde, 2% glutaraldehyde, and 0.1 M cacodylate buffer pH 7.4. TEM was carried out using standard techniques. Further details are described in the **Supplemental Methods**.

### Immunofluorescence studies

For immunofluorescence, the nephrocytes were dissected and fixed in 4% paraformaldehyde (PFA) in phosphate buffered saline (PBS) for 20 minutes. Subsequent antibody labeling was performed according to standard procedures. All primary antibodies used in this study are listed in **Suppl. Table 2**. Nuclei are marked by Hoechst 33342 (Sigma, B2261; 1:500-1:1000). Samples were imaged using a Zeiss LSM 980 applying Airyscan mode or an Abberior STEDYCON for Stimulated Emission Depletion (STED) microscopy.

## RESULTS

### *In-situ* visualization of the *Drosophila* filtration barrier by cryo-electron tomography

The *Drosophila* nephrocyte shapes membrane invaginations into an elaborate network termed labyrinthine channels. Entry into these channels is regulated by a filtration barrier comprising a basement membrane and a slit diaphragm (SD)^18,22,23^. The structural framework of the SD is established by Sns and Kirre, analogous to the function of their human orthologs nephrin and Neph1^22,23^ (**Figure 1A**). The SD architecture along the furrow-like labyrinthine channels features a fingerprint-like pattern in tangential sections (**Figure 1B**, **Supplementary Figure 1A-B**), corresponding to a regular, dot-like appearance in cross-sections (**Figure 3C**, **Supplementary Figure 1C**, three-dimensional visualization **Figure 1D**, **Supplementary Movie 1**). The SD proteins Sns and Kirre colocalize within this shared pattern (**Figure 1E-E’’**, **Supplementary Figure 1D-D’’**). To gain insight into the molecular architecture of the native *Drosophila* filtration barrier, we recorded cryo-electron tomograms (cryo-ET) of cryo-focused ion beam (FIB)-milled, on-grid lamellae from freshly isolated nephrocytes (n=6). The samples were kept in their native frozen hydrated state throughout all the experiments. All characteristic landmarks of the *Drosophila* filtration barrier, including the basement membrane, the labyrinthine channels on the cell surface, and the SDs, can be seen in three-dimensions (3D) (**Figure 1F-H**). The interior of the nephrocytes is filled with ribosomes, vesicles and various cytoskeletal elements (**Figure 1F-H**).

**Figure 1.**
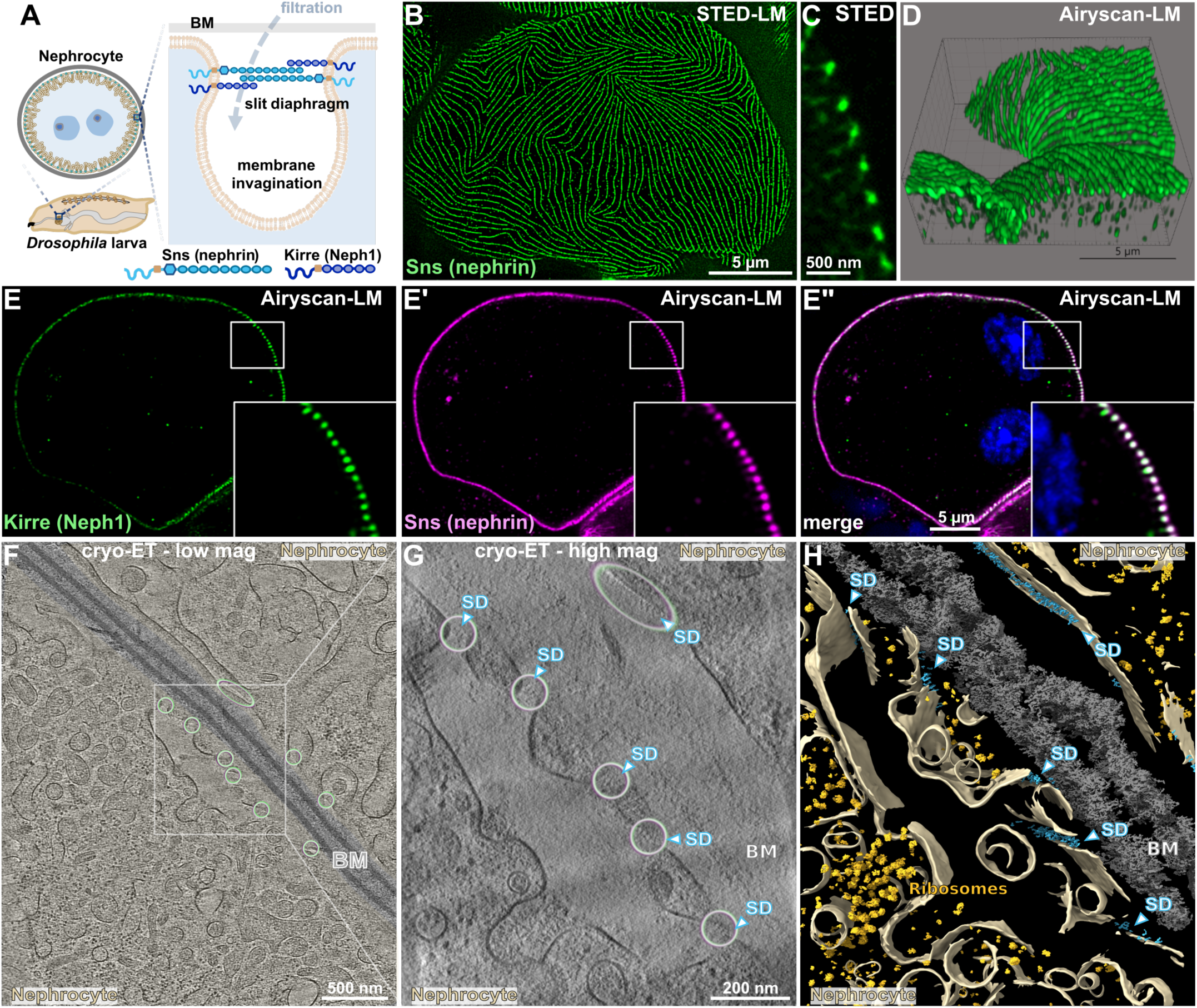
Comparative analysis of the nephrocyte slit diaphragm using light microscopy and cryo-electron tomography. (**A**)The schematic illustrates the anatomy and structure of nephrocytes including the SD, highlighting the two key SD proteins. (**B-C**) STED microscopy images of a nephrocyte expressing Myc-tagged Sns from a CRISPR-edited gene locus is shown. Myc staining reveals a fingerprint-like pattern of the tagged SD protein in a tangential section (B) and dot-like pattern in a cross-section (C). (**D**) Visualization using IMARIS software illustrates the relationship between linear and dot-like patterns based on Airyscan fluorescent microscopy of surface detail from nephrocytes stained for Myc-Sns. (**E-E’’**) Fluorescent microscopy of nephrocytes after co-labeling of (Myc)-Sns and Kirre shows colocalization of both SD proteins (enlarged inset). Nuclei are marked by Hoechst 33342 in blue. (**F**) 3 nm thick computational slice of a low-magnification tomogram showing the cross-section of two nephrocytes. The cytoplasm of the cells is gently highlighted in light yellow, the basement membrane in gray. In the cytoplasm, several vesicles and cytoskeletal elements can be seen. Encircled areas indicate regions containing SDs, where Sns and Kirre co-localize. (**G**) Computational slice of a tomographic reconstruction at higher magnification (1nm pixel size) revealing the nephrocyte SD (highlighted by blue arrows and circles). The SD appears textured, and not just as two lines connecting the plasma membranes. (**H**) Segmentation of the tomogram in (G), displaying the 3-dimensional architecture of the nephrocyte close to the cell surface, where nephrocyte SDs can be found. Light yellow: membranes, blue: nephrocyte slit diaphragms, gray: basement membrane, bright yellow: ribosomes.

### The slit diaphragm in *Drosophila* displays a fishnet-like pattern

In the tomograms, the SD can be visualized in 3D from all possible viewpoints (**Figure 2**, **Supplementary Movie 2**). Three characteristic views reveal: (i) a fishnet pattern in the ‘top view’ showing individual 52 nm long strands spanning the two plasma membranes at an angle of 50 degrees (**Figure 2B**), (ii) two parallel lines of “pearls on a string” density in the ‘membrane view’ (**Figure 2C**), and (iii) two electron dense parallel layers connecting the plasma membranes in the ‘classical view’, consistent with the micrographs from room temprature electron microscopy^22^ (**Figure 2D, Supplementary Figure 3**). The three views are orthogonal to each other and are shown in an isosurface visualization (**Figure 2H-K**). The SD has a width of approximately 44 nm (**Figure 3A**) in the extracellular space between two adjacent plasma membranes, which is similar to what has been previously described using RT-TEM with chemical fixation^22,23^. Due to a significantly enhanced resolution compared to RT-TEM (**Supplementary Figure 2A-H**), a periodic arrangement of the molecular densities composing the SD can be discerned already in the raw data (**Figure 1H-I**, **Figure 2B-D, Supplementary Movie 2**).

**Figure 2:**
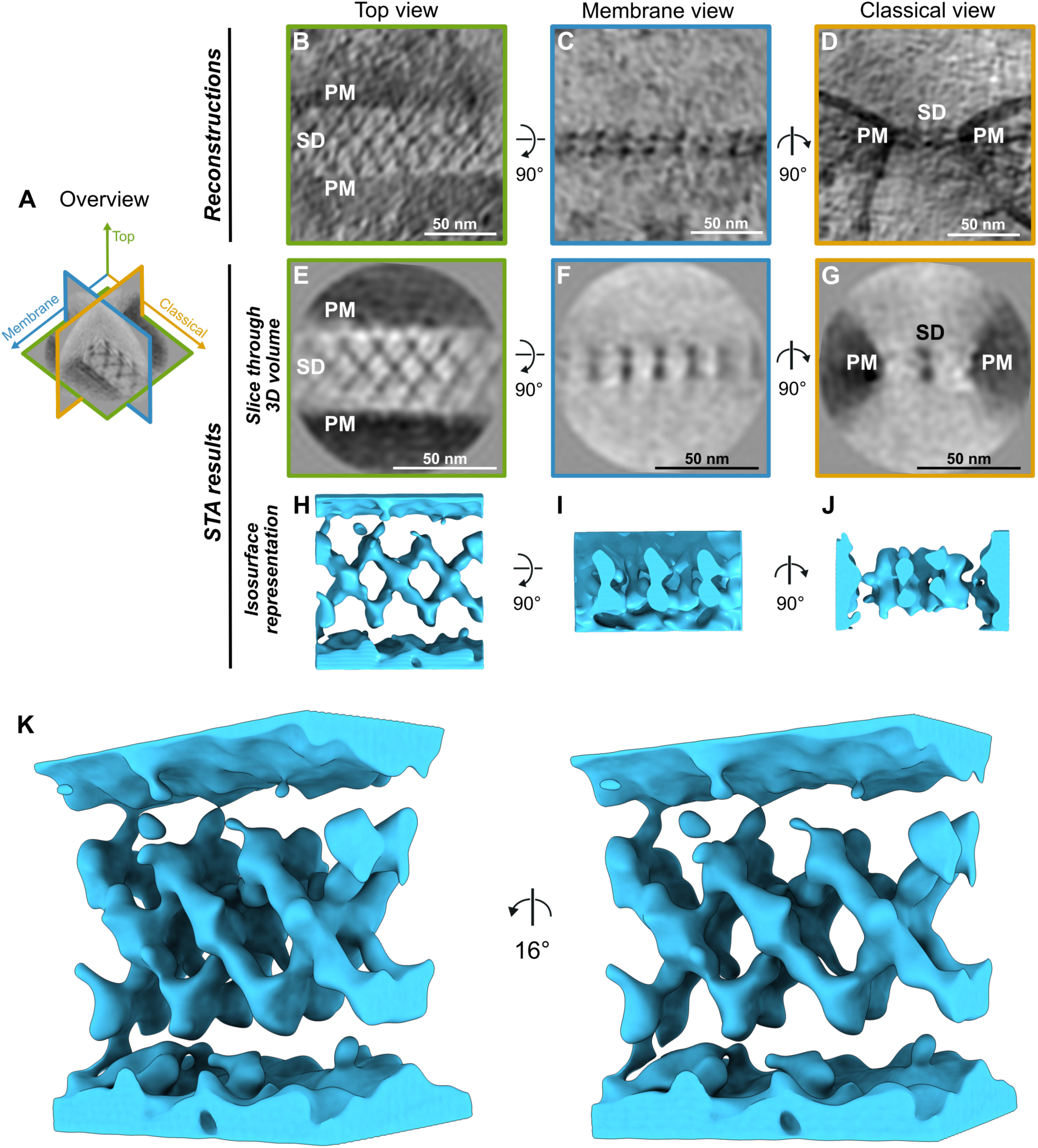
Cryo-ET reveals that the SD looks like a bi-layered fishnet. (**A**) Orthogonal planes through the SD average providing an overview of the 3D geometry. (**B-D**) 1 nm thick slices through the tomographic reconstructions showing the nephrocyte SD from three different perspectives ((B) top view revealing the fishnet-like pattern of the SD, (C) membrane view revealing two parallel lines of “pearls on a string”, (D) classical view displaying two parallel electron dense lines, reflecting the bi-layered architecture of the SD). (**E-G)** 1 nm thick slices through the 3D volume of the SD obtained after averaging, shown from the same perspectives as in B-D (pixel size: 2.7 Å). The slice showing the membrane view (F) illustrates the cross-section of the fishnet pattern at the center of the fishnet, parallel to the membrane. The slice showing the classical view (G) illustrates one of the cross-sections at the center of the fishnet perpendicular to the labyrinthine channel, and thus appears as two dots. (**H-J**) Isosurface representation of the cryo-ET map illustrating the 3-dimensional bi-layered fishnet-like architecture, shown from the same perspectives as in E-G (pixel size: 2.7 Å). Individual crossing strands forming the fishnet pattern can be identified. (**K**) Stereo-pair of the SD at an oblique angle showing the bi-layered architecture, where each of the layers resembles a fishnet.

To enhance the signal from inherently noisy cryo-ET data, subtomogram averaging is commonly used. This approach involves averaging the signal of many identical objects, thereby improving the signal-to-noise ratio (SNR). By averaging of 595 SD segments, a periodic arrangement of the molecular strands can be seen in higher contrast and improved resolution compared to the raw data in the individual tomograms (**Figure 2E-J**). The strands creating the fishnet pattern are approximately 4 nm thick and can be traced from one plasma membrane to the other. As views of the SD from several directions contributed to the average, the resulting cryo-ET map is isotropically resolved and depicts all SD views seen in the tomograms (**Figure 2**). When visualized in 3D, the overall architecture displays two fishnets stacked on top of each other (**Supplementary Movie 2**). Each fishnet is created by crossing strands. It can be sub-divided in triangular and rhombus shapes with a side length of 15 nm and angles of 110 and 70 degrees. Each strand of the rhombus pattern is ∼4 nm thick, resulting in structural holes of approximately 11 nm x 11 nm (**Figure 2E-J**, **Figure 3A**). These data indicate that the *Drosophila* SD consists of two identical layers, each resembling a fishnet, stacked on top of each other.

**Figure 3:**
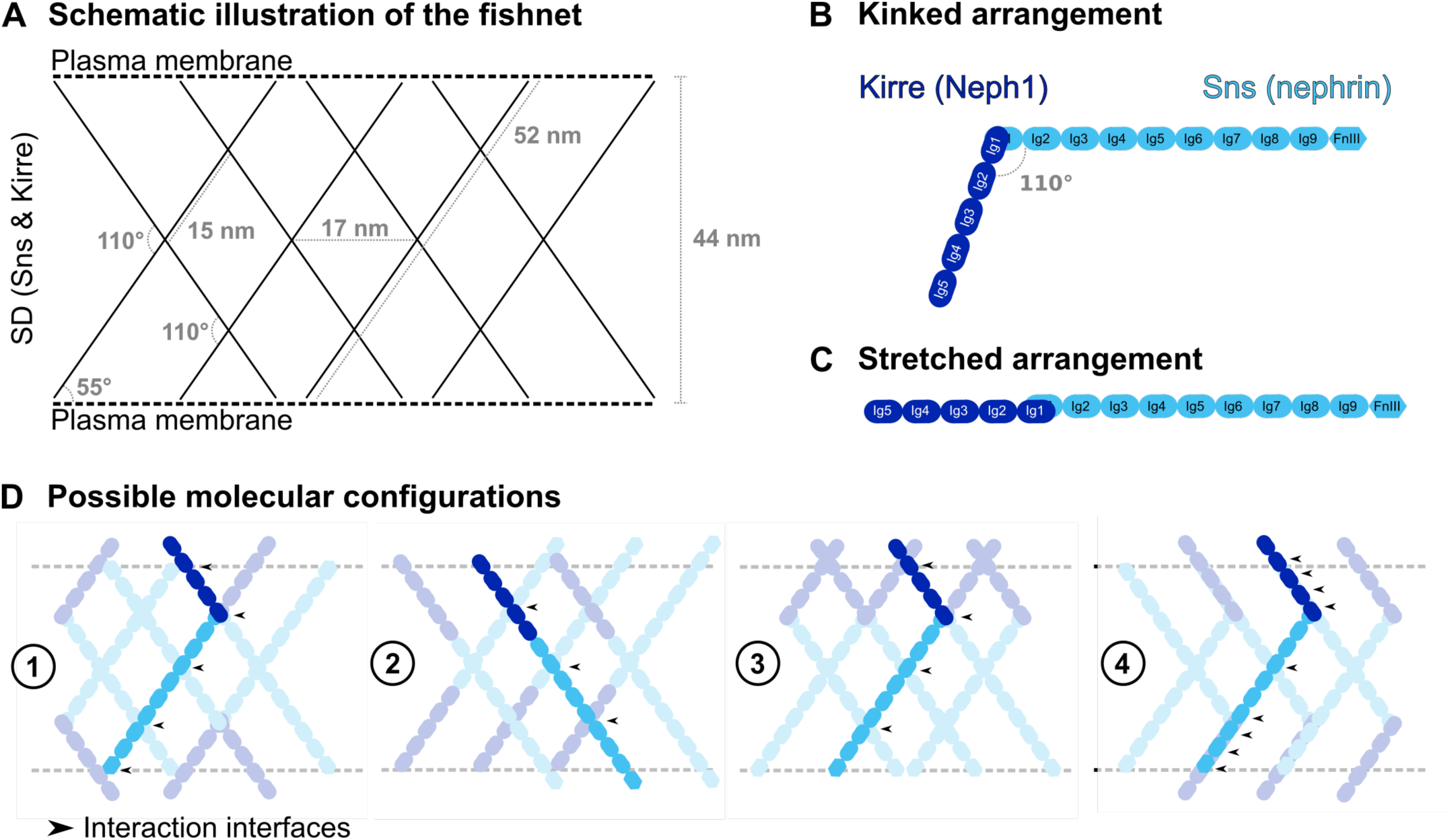
The possible configurations of Sns and Kirre within the SD. (**A**) Blueprint of the densities observed by cryo-ET. At the center of the SD, a prominent rhombus, with a 17 nm short diagonal and an interior angle of 110 degrees, can be seen. All relevant dimensions and spacings are indicated. (**B**) Kinked arrangement of Sns and Kirre, as observed in the X-ray structure by Özkan et al.^26^. (**C**) Stretched arrangement, where Sns and Kirre interact at an angle of 180°. (**D**) Possible molecular configurations based on the fishnet geometry of the SD. Arrows indicate potential interaction interfaces between crossing Ig domains.

### Proposed molecular configurations

Next, we populated the cryo-ET map of the SD with Sns and Kirre. Considering the dimensions of the SD (**Fig. 3A**) and the lengths of Kirre and Sns, one molecule alone is not long enough to span the distance between the two plasma membranes. Thus, Sns and Kirre proteins must interact in order to span the inter-membrane distance. Since the resolution is not sufficient to explicitly distinguish between Sns and Kirre, we constructed possible configurations that are structurally plausible (**Figure 3**). This resulted in four potential configurations for Sns and Kirre (**Figure 3D**). Configuration **1** is in accordance to the previous crystallographic studies and single particle analysis studies, where the Ig1 domains of Sns and Kirre interact at an angle between 90 to 110 degrees, in a kinked arrangement (**Figure 3B**)^26^. This is consistent with the angle of 110 degrees we measure in the rhombus of the fishnet pattern. In configuration **2**, Sns and Kirre interact at an angle of 180 degrees, in a stretched arrangement (**Figure 3C**). Although the interaction between the Ig1 domains of Sns and Kirre is predicted to be most stable at an angle of 90 to 110 degrees in solution, in their native environment a different structural arrangement could be imposed.

While configurations **3** and **4** are theoretically feasible, they appear biologically implausible. In configuration **3,** Kirre and Sns would derive exclusively from a single membrane side, and in configuration **4** there is a mismatch with the observed cryo-ET map – either due to the absence of molecules in regions where there is a density, or due to the overlap of multiple Ig domains resulting in thicker strands than observed. Each suggested configuration generates specific potential interaction interfaces, where individual Ig domains are found in close proximity to each other and may interact (**Table 1**).

**Table 1:** Interaction interfaces between Sns and Kirre for each proposed configuration.

|  |  |  |
| --- | --- | --- |
| **Model 1** | Heterophilic interactions | Kirre-Ig1 and Sns-Ig1;<br>Kirre-Ig4 and Sns-FnIII-like |
|  | Homophilic interactions | Sns-Ig4 and Sns-Ig4;<br>Sns-Ig7/Ig8 and Sns-Ig7/Ig8 |
| **Model 2** | Heterophilic interactions | Kirre-Ig1 and Sns-Ig1;<br>Kirre-Ig2/3 and Sns-Ig6/7;<br>Kirre-Ig5 and Sns-Ig9/FnIII-like |
|  | Homophilic interactions | Sns-Ig3 and Sns-Ig3 |
| **Model 3** | Heterophilic interactions | Kirre-Ig1 and Sns-Ig1 |
|  | Homophilic interactions | Kirre-Ig4/5 and Kirre-Ig4/5;<br>Sns-Ig4 and Sns-Ig4;<br>Sns-Ig7/Ig8 and Sns-Ig7/Ig8 |
| **Model 4** | Heterophilic interactions | Kirre-Ig1 and Sns-Ig1;<br>Kirre-Igs1-5 and Sns-Igs7-FnIII-like |
|  | Homophilic interactions | Sns-Ig4 and Sns-Ig4 |

### The fishnet-like pattern is abolished upon *sns* silencing and shifts into the labyrinthine channels upon mistrafficking associated with *Rab5*-RNAi

To examine whether the fishnet pattern requires the presence of the slit diaphragm protein Sns, we performed cryo-ET in samples from animals expressing *sns*-RNAi. The typical Sns-derived signal in light microscopy disappeared upon knockdown of the *Drosophila* nephrin ortholog (**Supplementary Figure 3A-B’’**), confirming efficient silencing. In absence of its binding partner, the linear pattern of Kirre was replaced by a finely dotted pattern in clusters (**Figure 4A**), while the mislocalized Kirre largely adhered to the surface (**Figure 4B**). The altered configuration confirms that the Neph1 ortholog alone is insufficient for slit diaphragms formation and consequently, the characteristic labyrinthine channels were no longer formed (**Figure 4C-D**). In RT-TEM and cryo-ET acquisitions, we observed electron-dense clusters decorating the cell membrane (**Figure 4C-F**), but the SDs with their fishnet-like pattern were abrogated. Consistent with previous findings^22,23^, we conclude that the presence of Sns is essential for the formation of the nephrocyte SD, including its newly discovered fishnet-like pattern.

**Figure 4:**
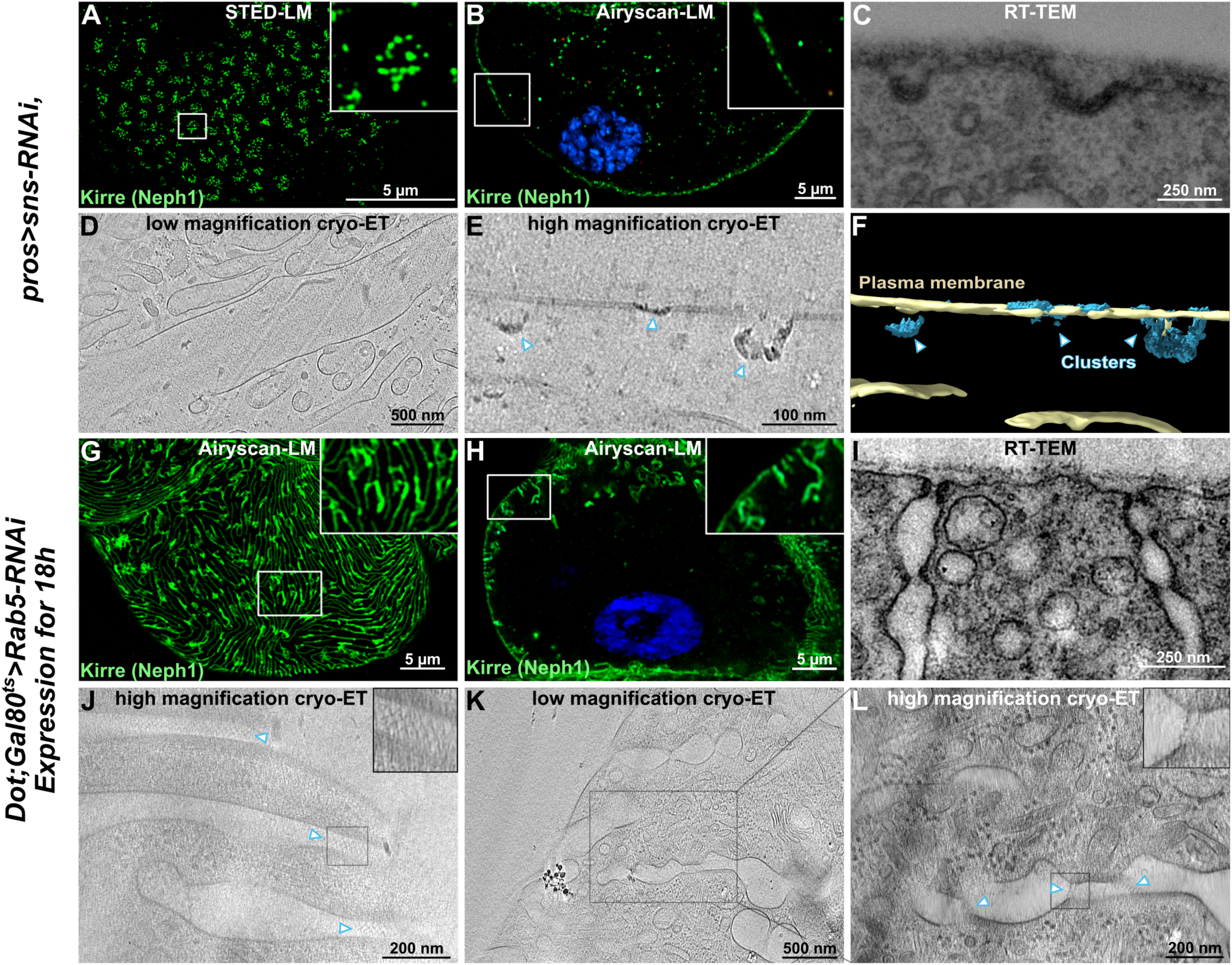
The fishnet-like pattern is absent with *sns* silencing and appears ectopically upon *Rab5*-associated disruption of *sns* trafficking. (**A-B**) Fluorescence microscopy of nephrocytes expressing *sns*-RNAi reveals a finely dotted pattern of Kirre in tangential sections (A) while the SD protein largely maintains association with the cell surface indicated by cross-sections (B). (**C**) RT-TEM image shows detail of a nephrocyte expressing *sns*-RNAi. In absence of the SD protein, SD formation is abrogated while electron-dense pits are formed on the surface. (**D**) Slice through low-magnification tomographic reconstruction of the cross-section through two *sns* KD nephrocytes. No channels or invaginations can be identified on the cell surface. (**E**) Section through high-magnification tomographic reconstruction revealing electron dense clusters close to the outer membrane of the nephrocyte. (**F**) Segmentation illustrates the 3D geometry of the clusters inside the plasma membrane of the nephrocyte (blue: clusters, light yellow: plasma membrane). (**G-H**) Fluorescence microscopy of nephrocytes after short-term expression of *Rab5*-RNAi for 18 h shows a partially maintained SD pattern in tangential sections (G) as well as ectopic Sns protein protruding in lines from the nephrocyte surface in cross-sections (H). (**I**) RT-TEM image of two labyrinthine channels of a nephrocyte expressing *Rab5*-RNAi for 18 h reveals ectopic SD formation. (**J**) Slice through high-magnification reconstruction after cryo-ET of nephrocyte expressing *Rab5-*RNAi showing the labyrinthine channels from the top. Insert: Zoom-in revealing the fishnet-pattern, similar to the one found in the wild-type nephrocyte. (**K**) Slice through low-magnification tomographic reconstruction, where SDs can be found translocated towards the interior of the cell. (**L**) Section of higher magnification tomographic reconstruction of the same region of interest as in (K). Insert: Zoom-in revealing classical views of the SDs found in the nephrocyte expressing *Rab5*-RNAi.

Finally, we investigated a genetic background that exhibits mislocalized slit diaphragms without loss of directly SD-associated proteins. Knockdown of *Rab5*, a small GTPase involved in regulation of endocytosis and cargo sorting, is known to disrupt Sns trafficking and leading to ectopic formation of SD within the network of labyrinthine channels^19^. After short-term *Rab5* silencing for 18 h, we detected the slit diaphragm protein in a mildly rarefied and blurred linear pattern in nephrocyte tangential sections (**Figure 4G**). Cross-section now additionally reveals Sns protein protruding from the surface (**Figure 4H**). This suggests presence of ectopic SD protein within the channel network, and slit formation was confirmed by RT-TEM (**Figure 4I**). The SD bi-layered fishnet pattern was maintained on the surface, as revealed by cryo-ET (**Figure 4-J**). With a spacing of approximately 17 nm measured between protein strands and a width of roughly 47 nm measured from membrane to membrane, the fishnet pattern corresponds to what was found for wild-type nephrocytes. Simultaneously, we observed the presence of SDs translocated from the cell surface deeper into the channel network (**Figure 4-K-L**), consistent with observations by light microscopy and RT-TEM (**Figure 4-G-I**). Thus, the disruption of Sns trafficking after *Rab5* silencing results in ectopic appearance of the fishnet pattern. However, short-term silencing of Rab5 seems to affect the localization of the SDs, but not their assembly into a fishnet pattern.

## DISCUSSION

The *Drosophila* nephrocyte SD features a bi-layered, fishnet-like pattern, which is absent following knockdown of *sns*, the *Drosophila* nephrin ortholog, and appears ectopic after SD mistrafficking. The structure of the *Drosophila* SD is remarkably similar to the murine SD, as both resemble a fishnet, have a practically identical strand thickness, and the strands spanning the space between the two plasma membranes form rhombus shapes. Thus, the overall molecular architecture is very similar. There are a few differences: (i) the spacing between the two plasma membranes is 44 nm, compared to 53 nm in the murine SD. (ii) The spacing between the strands spanning the two plasma membranes is larger, i.e. 17 nm compared to 11nm in the murine SD. In combination with the smaller plasma membrane spacing, only one rhombus shape appears in the center of the SD compared to two rhombus shapes in the murine SD. (iii) The nephrocyte SD appeared in all our images as a double layer, in contrast to a single layer or multiple layers in the murine SD. Image analysis and subtomogram averaging indicated that the distance between the two layers remains stable, and that the two layers are potentially connected to each other. (iv) The width of the SD appears very stable, in stark contrast to the murine SD leading to a better resolved cryo-ET map.

The remarkable similarity of the SD’s architecture between *Drosophila* and mammals underscores the configuration’s functional significance. The fishnet pattern seems plausibly linked to the demanding functional requirements of the slit diaphragm. A fishnet provides strong mechanical stability while enabling size-selective permeability. Recent studies indicated that the SD is a highly dynamic structure, undergoing rapid cycles of endocytosis and recycling^19^. Since specific interaction interfaces between the immunoglobulin domains seem to define the molecular configuration, each layer may provide the template to guide newly delivered components of the opposing sheet during constant turnover. This molecular backup is fit to explain how continuous renewal of the SD occurs without protein leakage, despite constant filtration. A maximal measured mesh size of 17 nm just allows the passage of albumin molecules, which directly corresponds to observations from tracer studies^24^. The wider mesh size compared to mice might contribute to the marginally higher size cut-off of the nephrocyte, which allows the passage of albumin but not transferrin^24^.

Previous studies involving staining with heavy metals could not achieve the resolution necessary to clearly discern individual strands. In cryo-ET, the sample is just frozen to liquid nitrogen temperatures without any subsequent treatment to enhance the contrast. With the technology we use here -cryo-electron tomography of FIB-milled plunge frozen samples-even atomic resolution has been achieved within cells ^27^.

Different models have been proposed for the mammalian SD^14–17^. Single particle electron microscopy and crystallography of SYG-1 and SYG-2, orthologs of nephrin (Sns) and Neph1 (Kirre), predicted an interaction angle of 90-110 degrees between the Ig1-domains of these proteins^26^. Intriguingly, this is the angle that we obtained between the crossing strands that define the fishnet pattern. Assuming this 110 degree interface, with both proteins being expressed from both sides of the channel and no molecular clashes, only model **1** for the configuration of Sns (nephrin) and Kirre (Neph1) appears plausible within the observed fishnet-like pattern (**Figure 3-D, model (1)**). If the interaction angles between the molecules are arbitrary or the molecules are bent, several other models become possible. Each of the proposed configurations present potential interaction interfaces between the Ig domains of Sns and Kirre that may facilitate the formation of the fishnet pattern. While it is known that Ig4, Ig6 and the FnIII-like domain of Sns are crucial for the formation of the SD^23^, these characterized interactions do not unambiguously verify or exclude any of the here presented possible molecular configurations. Therefore, gaining a deeper understanding of potential interactions and potential affinity between the respective Ig domains is crucial in order to understand the arrangement of the proteins within the SD. Future mutation studies may also aid in verifying the proposed configurations.

Finally, our findings emphasize the relevance of *Drosophila* nephrocytes as a valuable podocyte model for studying slit diaphragm biology — addressing the gap left by lack of this structure from *in vitro* models for mechanistic studies and drug discovery.

Taken together, our findings show that the SD has a fishnet-like architecture and emphasize the conservation of this distinct pattern among species, underscoring its biological significance.

## DISCLOSURE STATEMENT

The authors declare no competing interests.

## AUTHOR CONTRIBUTIONS

D.M., K.L., A.N.B., and M.H.: Prepared samples for cryo-electron tomography (cryo-ET). D.M.: Focused ion beam (FIB)-milled lamellae, acquired cryo-ET data and processed the data. K.L.: Reared the flies, prepared samples for light microscopy, acquired confocal and STED data. D.M., K.L., A.N.B., and T.H. Prepared the data figures M.P.S.: Performed the processing and refined the final density map. T.H. and M.H.: Prepared samples for room temperature transmission electron microscopy (RT-TEM), and acquired RT-TEM data. M.P.S.: Designed the research. A.S.F. and T.H.: Designed and supervised the research. D.M., K.L., T.H., and A.S.F.: Wrote the manuscript, with contributions from all authors.

## ACKNOWLEDGEMENTS

We are grateful to Gerd Walz for valuable advice. We thank C. Meyer for outstanding technical support. We are grateful for support by F. Ditengou, Lighthouse Core facility, University of Freiburg, regarding confocal microscopy. Lighthouse Core Facility is funded in part by the Medical Faculty, University of Freiburg (Project Numbers 2021/A2-Fol; 2021/B3-Fol) and the DFG (Project Number 450392965). We thank the Frankfurt Center for Electron Microscopy for measurement time. We thank Maria Ericsson (Harvard Medical School Electron Microscopy Facility) for technical assistance. We thank the Bloomington Drosophila Stock Center for providing fly stocks.

The Core Facility for Electron Microscopy (EMcore) at the University Freiburg Medical Center— IMITATE is registered with the DFG (German Research Foundation) under the reference number

## Data availability statement

No large-scale data sets from high-throughput analyses were generated or analyzed in this study, with the exception of whole exome sequencing data.

Unprocessed confocal or electron microscopy images are available upon reasonable request. All remaining data are included within the manuscript.

## Notes

**Funding:** The Deutsche Forschungsgemeinschaft funded D. M. and A.N.B. (Research Training Group iMOL / GRK 2566/1), M.P.S. (FR 1653/14-1), T. H. (HE 7456/7-1 and project-ID 431984000 – SFB 1453), F.G. (CRC 1192, GR3933/1-1). T. H. acknowledges support from the Heisenberg Programme of the DFG (HE 7456/6-1)

### Competing Interest Statement

The authors have declared no competing interest.

