## Supplemental_Data for "The slit diaphragm in *Drosophila* features a bi-layered, fishnet-like architecture"

<u>Table of contents - Supplementary Information</u>

**Supplemental Methods**

**Supplementary Figure 1: Additional imaging illustrating the nephrocyte filtration barrier.**

**Supplementary Figure 2: Comparison of the resolution in nephrocytes between conventional room temperature transmission electron microscopy (RT-TEM) and cryo-electron tomography (cryo-ET).**

**Supplementary Figure 3: Confirmation of effective *sns* silencing by immunofluorescence.**

**Supplementary Table 1: Additional fly strains.**

**Supplementary Table 2: Primary antibodies.**

**Supplementary References.**

### Supplemental Methods

#### Cryo-electron tomography.

***Sample preparation***

For cryo-electron tomography, nephrocytes were dissected from *Drosophila* L3 larvae in Schneider's medium containing 10% glycerol [1]. They were then centrifuged (700 xg, 8 min) onto poly-L-lysin coated EM grids (Plano, copper/palladium, G2019D 100 mesh, G2018 75 mesh, or G2050C 50 mesh). The EM grids were coated with Formvar to serve as a support film, and subsequently coated with poly-L-lysine (PLL) to facilitate adhesion of the nephrocytes. PLL coating was performed by glow-discharging the grids for 15 s, applying 8 μl of 0.1% (w/v) PLL in $H_2O$ (Sigma-Aldrich, St. Louis, Missouri, US), incubating for 20 min and washing them for 5 s in MilliQ® water [2-4]. After centrifugation, EM grids with nephrocytes were manually blotted (VWR, Qualitative filter paper, 413) and plunge frozen in liquid ethane (Vitrobot Mark IV, Thermo Fischer Scientific). The Vitrobot chamber was kept at 20°C and 100% relative humidity. Samples were stored at -196˚C in liquid nitrogen.

***Cryogenic confocal laser scanning microscopy***

To facilitate sample targeting in the cryo focused ion beam (FIB)-milling step, the plunge frozen grids were imaged at -195°C by confocal laser scanning microscopy (cryo-CLSM) (CMS196 cryo-stage, Linkam, Salfords, United Kingdom; LSM700, Carl Zeiss, Jena, Germany). Optical configurations were adjusted to capture the autofluorescence of the nephrocytes in the green channel, and the reflection from the grids in the far-red channel with excitation wavelengths of 488 and 639 nm, respectively. Images were acquired with a 5x/ NA 0.16 objective. Data acquisition was performed with Zeiss ZEN 2009 (blue) v2.1.

#### *Cryo FIB-milling*

After cryogenic confocal laser scanning microscopy, the plunge frozen grids were clipped into FIB autogrids (#1205101, Thermo Fisher Scientific Inc.). On-grid lamellae were produced by FIB-milling in a Helios Nanolab 600i (Thermo Fisher Scientific Inc), as described previously [5] [6].

#### *Cryo-ET data acquisition*

Tilt series were acquired with SerialEM v4.1.0 beta [7] in a Titan Krios transmission electron microscope (Thermo Fisher Scientific Inc.) operated at 300 keV in nanoprobe EFTEM mode, equipped with a GIF Quantum S.E. post-column energy filter in zero-loss peak mode and a K3 Summit detector (Gatan, Pleasanton, USA). Tilt series were recorded at a nominal magnification of 33 000x (1.34 Å per pixel) in super-resolution and dose-fractionation modes. A total of 52 images was acquired in dose-symmetric scheme using a tilt increment of 2°, starting from the pre-tilted lamella. Stage tilt angles ranged from -66° to +36°. The total dose per micrograph was at 3.125 $e^-/Å^2$ (total dose for 52 images: 162,5 $e^-/Å^2$), and the defocus was set to -5 $\mu$m.

#### *Cryo-ET data processing*

Tilt series were processed and subtomogram averaging was performed using RELION-5[8]. Movies were motion corrected using the MotionCorr2 wrapper [9] and CTF estimation was performed with ctffind-4.1.14 [10]. Alignment was performed using the AreTomo2 wrapper or, alternatively, patch tracking in IMOD [11]. Tomograms were reconstructed (binned pixel size 10.72 Å) and particles were manually selected and pre-oriented using ArtiaX in UCSF ChimeraX [12] [13]. 848 sub-tomograms (box size 96, pixel size 10.72 Å) containing the two plasma membranes and the extracellular space with the SD were used for an initial 3D classification to select the best particles, yielding a list of 595 particles. Subsequent rounds of 3D refinement and gradual unbinning to a pixel size of 2.68 Å resulted in the final map at 39.58 Å. For **Figure 1G**, the tomogram was reconstructed using IMOD's SIRT-like reconstruction. For **Figures 2B-D**, the tomograms were processed with tom_deconv[14] and IsoNet[15]. For **Figures 4D-E**, the tomograms were reconstructed using IMOD'S SIRT-like reconstruction. For **Figures 4J-L**, tomograms were denoised with the cryoCARE wrapper[16] within RELION-5, and subsequently processed with IsoNet. Segmentations were generated with Dragonfly software, Version 2024.1, for Windows [17].

### Supplementary Figures

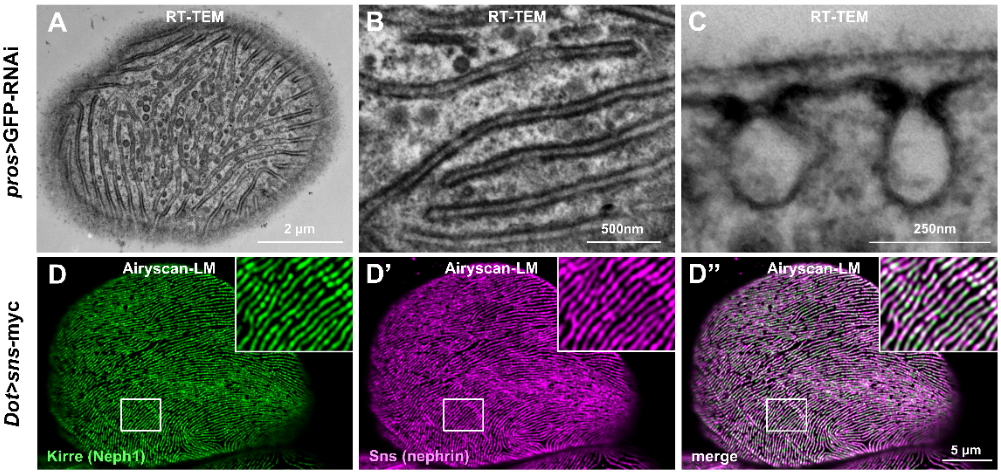


**Supplementary Figure 1: Additional imaging illustrating the nephrocyte filtration barrier.**

**(A-C**) RT-TEM images of control nephrocytes are shown. (A-B) Tangential sections illustrate the fingerprint-like pattern formed by the slit diaphragm (A), which are discernible by parallel electron-dense lines with higher magnification (B). Magnified RT-TEM image of a nephrocyte surface detail in a cross-section shows slit diaphragms as thin lines between electron-dense areas on top of the oval membrane invagination.

**(D-D")** Immunofluorescence microscopy image acquired in Airyscan mode shows the colocalization of slit diaphragm proteins Sns (ortholog of nephrin) and Kirre (ortholog of Neph1) in a fingerprint-like pattern.

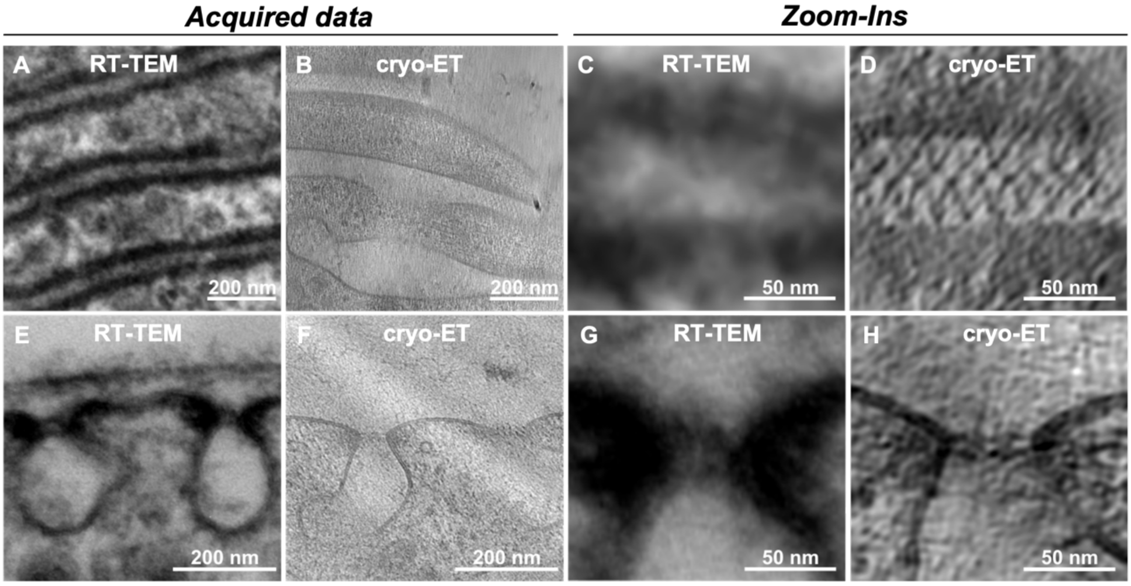


**Supplementary Figure 2: Comparison of the resolution in nephrocytes between conventional room temperature transmission electron microscopy (RT-TEM) and cryo-electron tomography (cryo-ET).**

**(A, E)** RT-TEM overview images of few nephrocyte slit diaphragms shown in tangential and in the cross-section, respectively.

**(B,F)** Slices of the tomographic reconstructions obtained by cryo-ET for the tangential section and cross-section across the labyrinth channels, at the same magnification as the RT-TEM images shown for comparison. The bi-layered (F), fishnet-like (B) SD can be seen in the extracellular space within the channels. A richer image can be appreciated, displaying individual ribosomes, cytoskeletal fibers, and many cellular features that are absent in the RT-TEM images.

**(C,G**) Zoom-ins of RT-TEM images of a labyrinth channel shown in tangential (C) and in the cross-section (G).

**(D,H**) Zoom-ins of the tomographic reconstructions of the cryo-ET dataset, revealing the fishnet-like arrangement (D) and the bi-layered architecture (H) of the SD, which is not discernible in the resolution that is possible using RT-TEM.

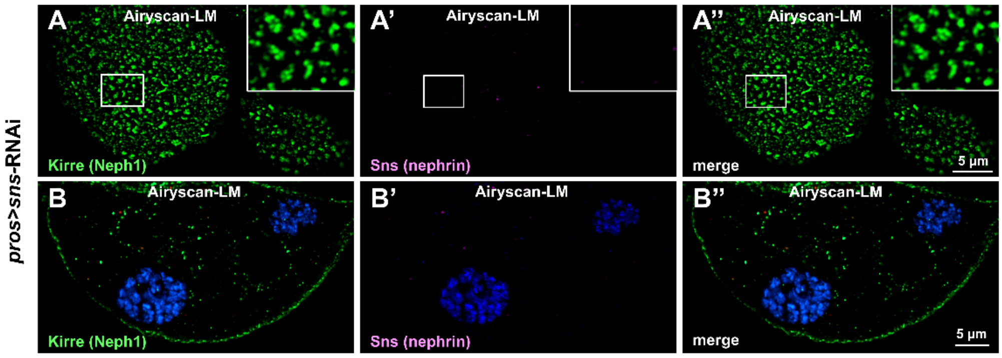


**Supplementary Figure 3: Confirmation of effective *sns* silencing by immunofluorescence.**

**(A-B")** Fluorescence microscopy images acquired in Airyscan mode of nephrocytes after silencing of *sns* in tangential (A-A") and cross-section (B-B") illustrate the absence of Sns (nephrin) protein. The slit diaphragm protein Kirre (Neph1) exhibits round clusters (A) instead of the normal linear pattern without its binding partner.

| Fly strain | Source |
|---|---|
| tubP-GAL80$^{ts}$ | Bloomington Drosophila Stock Center #7019 |
| UAS-EGFP-RNAi | Bloomington Drosophila Stock Center #41553 |
| Dorothy-*GAL4* | Bloomington Drosophila Stock Center #6903 |
| UAS-*Rab5*-RNAi | Bloomington Drosophila Stock Center #34832 |
| Myc-Sns | Lang et al., 2022; PMID: 35876643 |

**Supplementary Table 1:** Additional fly strains

| Antibody | Source |
|---|---|
| mouse anti-Myc | Cell Signaling Technologies #2276 |
| rabbit anti-Kirre | custom antibody (Davids Biotechnologie) using peptides HAKSKKNKSSQSSHHGDSS and EEHHLPEGVRAALIIRDSKAT |

**Supplementary Table 2:** Primary antibodies

### SUPPL. REFERENCES


1. Bäuerlein FJ, Renner M, Chami DE, Lehnart SE, Pastor-Pareja JC, Fernández-Busnadiego R. Cryo-electron tomography of large biological specimens vitrified by plunge freezing. *BioRxiv*. 2021:2021.2004. 2014.437159.
2. Mazia D, Schatten G, Sale W. Adhesion of cells to surfaces coated with polylysine. Applications to electron microscopy. *J Cell Biol*. Jul 1975;66(1):198-200. doi:10.1083/jcb.66.1.198
3. Harapin J. *Structural characterization of macromolecular complexes within thick specimens using cryo-focused-ion-beam scanning electron microscopy (cryo-FIB-SEM) and cryo-electron-tomography (cryo-ET)*. University of Zurich; 2017.
4. Shahmoradian SH, Galiano MR, Wu C, et al. Preparation of primary neurons for visualizing neurites in a frozen-hydrated state using cryo-electron tomography. *J Vis Exp*. Feb 12 2014;(84):e50783. doi:10.3791/50783
5. Kelley K, Raczkowski AM, Klykov O, et al. Waffle Method: A general and flexible approach for improving throughput in FIB-milling. *Nat Commun*. Apr 6 2022;13(1):1857. doi:10.1038/s41467-022-29501-3
6. Birtasu AN, Wieland K, Ermel UH, et al. The molecular architecture of the kidney slit diaphragm. *BioRxiv*. 2023:2023.2010. 2027.564405.
7. Mastronarde DN. SerialEM: A Program for Automated Tilt Series Acquisition on Tecnai Microscopes Using Prediction of Specimen Position. *Microscopy and Microanalysis*. 2003;9(S02):1182-1183. doi:10.1017/s1431927603445911
8. Burt A, Toader B, Warshamanage R, et al. An image processing pipeline for electron cryo-tomography in RELION-5. *FEBS Open Bio*. Nov 2024;14(11):1788-1804. doi:10.1002/2211-5463.13873
9. Zheng SQ, Palovcak E, Armache JP, Verba KA, Cheng Y, Agard DA. MotionCor2: anisotropic correction of beam-induced motion for improved cryo-electron microscopy. *Nat Methods*. Apr 2017;14(4):331-332. doi:10.1038/nmeth.4193
10. Rohou A, Grigorieff N. CTFFIND4: Fast and accurate defocus estimation from electron micrographs. *J Struct Biol*. Nov 2015;192(2):216-221. doi:10.1016/j.jsb.2015.08.008
11. Mastronarde DN, Held SR. Automated tilt series alignment and tomographic reconstruction in IMOD. *J Struct Biol*. Feb 2017;197(2):102-113. doi:10.1016/j.jsb.2016.07.011
12. Ermel UH, Arghittu SM, Frangakis AS. ArtiaX: An electron tomography toolbox for the interactive handling of sub-tomograms in UCSF ChimeraX. *Protein Sci*. Dec 2022;31(12):e4472. doi:10.1002/pro.4472
13. Meng EC, Goddard TD, Pettersen EF, et al. UCSF ChimeraX: Tools for structure building and analysis. *Protein Sci*. Nov 2023;32(11):e4792. doi:10.1002/pro.4792
14. Tegunov D, Cramer P. Real-time cryo-electron microscopy data preprocessing with Warp. *Nat Methods*. Nov 2019;16(11):1146-1152. doi:10.1038/s41592-019-0580-y
15. Liu YT, Zhang H, Wang H, Tao CL, Bi GQ, Zhou ZH. Isotropic reconstruction for electron tomography with deep learning. *Nat Commun*. Oct 29 2022;13(1):6482. doi:10.1038/s41467-022-33957-8

16. Buchholz T-O, Krull A, Shahidi R, Pigino G, Jékely G, Jug F. Content-aware image restoration for electron microscopy. *Methods in cell biology*. 2019;152:277-289.
17. *Dragonfly*. Version 2024.1. Comet Technologies Camada Inc.; https://www.theobjects.com/dragonfly
